# Spermidine Is The Main Polyamine Required By Intracellular Parasites For Survival Within Host Erythrocytes

**DOI:** 10.1101/2023.02.01.526700

**Authors:** Pallavi Singh, Choukri Ben Mamoun

**Affiliations:** Department of Internal Medicine, Section of Infectious Diseases, Yale School of Medicine, New Haven, Connecticut, USA

**Keywords:** *Babesia duncani*, *Plasmodium falciparum*, erythrocytes, polyamine, putrescine, spermidine, hypusination

## Abstract

Intracellular eukaryotic pathogens such as *Babesia* and *Plasmodium*, the agents of human babesiosis and malaria, require salvage or de novo synthesis of several nutrients for survival in human erythrocytes. One such nutrient is putrescine, which is either transported from the host or synthesized from ornithine and serves as a precursor for the biosynthesis of two other polyamines: spermidine and spermine. However, the specific polyamines required by these parasites for survival and the molecular process they control remain unknown. We show in both *B. duncani* and *P. falciparum* that spermidine is the main product of the polyamine biosynthesis machinery required for parasite survival. Simultaneous inhibition of spermidine synthesis from putrescine and catabolism from spermine results in cell death and parasite survival can only be rescued by spermidine. Finally, we demonstrate that spermidine’s essential function in these parasites is through regulation of protein translation via hypusination of the translation initiation factor eIF5A.

## Introduction

Apicomplexan protozoans comprise a broad group of obligate intracellular parasites that cause various diseases in humans and animals with major economic impact worldwide. Human malaria (*Plasmodium* species) and human babesiosis (*Babesia* sp.) are two important diseases caused by apicomplexan parasites with a significant clinical relevance and a global health impact. While malaria is mosquito-borne and babesiosis is tick-borne, clinical presentations in both human malaria and human babesiosis are similar and include relapsing fever, chills, sweats, headaches, nausea, and hemolytic anemia.

The majority of human babesiosis clinical cases worldwide have been linked to *B. microti*, which is transmitted by *Ixodes* ticks and carries the smallest genome among Apicomplexa ^1, 2^. Other babesiosis cases in Europe and United States have been linked to other *Babesia* species including *B. divergens, B. MO1, B. venatorum, B. duncani* and *B. odocoilei* ^3–5^. A crucial but poorly understood factor in understanding *Babesia* pathogenesis and virulence is how these parasites survive within host erythrocytes. The recent development of the *B. duncani* In culture-In mouse (ICIM) propagation model has enabled investigation of the molecular mechanisms underlying survival of *Babesia* parasites within erythrocytes and their interaction with the host ^6^. The model combines continuous in vitro culture of the parasite in human red blood cells with an optimized in vivo model of lethal infection in C3H-HeJ mice ^6^. The intraerythrocytic life cycle of *B. duncani* initiates following invasion of a human erythrocyte by a free merozoite. Upon invasion, the parasite develops into metabolically active forms (ring and filamentous stages) and multiplies to produce four daughter merozoites within the infected cell ^6^. Following rupture of the host cell, the released merozoites initiate new intraerythrocytic cycle leading to rapid increase in parasite burden. As in malaria, the repeated rounds of parasite development and multiplication and destruction of host erythrocytes are responsible for all the clinical symptoms associated with human babesiosis.

Survival of apicomplexan parasites in host RBCs is absolutely dependent on the uptake and/or de novo synthesis of several essential nutrients ^7–9^. Accordingly, gene conservation or loss and in some instances, gene redundancy have been crucial in determining the reliance of these parasites on either the uptake of exogenous nutrients from host plasma or their synthesis de novo from available precursors. Among these nutrients are the polyamines putrescine, spermidine and spermine, which have been shown to play critical roles in the viability of intraerythrocytic parasites by regulating cell growth and differentiation ^10^. While *Plasmodium* species use the de novo pathway of polyamine biosynthesis from L-arginine ^11,12^, the pathway used by *Babesia* remains unknown. Equally important, in neither organism have the specific polyamines required for survival and the cellular processes controlled by such polyamines been elucidated.

Here we provide evidence that *B. duncani* lacks a de novo polyamine biosynthesis pathway and instead relies on the uptake of exogenous putrescine from the host for survival within human erythrocytes and operation of critical cellular activities, chief among them is protein translation. Furthermore, we demonstrate that spermidine synthesized from salvaged putrescine or spermine is the key polyamine required for parasite proliferation and survival in human RBCs. Consistent with the modulation of *B. duncani* protein translation by polyamines, pharmacological studies showed that inhibition of hypusination of the parasite translation initiation factor 5A (eIF5A) from spermidine results in parasite death. The finding that spermidine is the key polyamine for intraerythrocytic parasitism was further confirmed in the human malaria parasite *P. falciparum*, which uses a de novo pathway for polyamine biosynthesis.

## Results

### Putrescine is essential for survival of *B. duncani* in human erythrocytes

Recent studies have shown successful continuous propagation of *B. duncani* in human erythrocytes in DMEM/F12 but not the base DMEM culture media ^13^ (Fig. 1A). Parasites maintained in DMEM/F12 over multiple 3-day propagation cycles showed increased parasite load overtime, with parasitemia doubling every 22-24 h, whereas those transferred from DMEM/F12 to the DMEM base medium experienced a steady decline in parasite survival over time with no viable parasites detected by the 5^th^ cycle of growth in this medium (Fig. 1A). Microscopy analysis of the morphology of parasites following transfer to DMEM medium showed accumulation of highly stressed and dead parasites displaying condensed chromatin as well as abnormal cellular morphology (Fig. 1B). Examination of the nutritional composition of the two media identified 15 components that are present in DMEM/F12 but not in DMEM media (Table 1). These include six amino acids (Ala, Asp, Asn, Cys, Glu and Pro), two vitamins (biotin and vitamin B12), four salts (cupric sulfate, ferric sulfate, magnesium chloride, zinc sulfate), two lipids (linoleic acid (LNA) and lipoic acid (LA)) and one polyamine (putrescine). To investigate which of the components is vital for the growth of the parasite, *B. duncani* cultures were grown in DMEM medium either alone or supplemented with the missing amino acids, vitamins, inorganic salts, lipids (LNA+LA) or putrescine or various combinations of these nutrients. Parasite growth was monitored for 15 days with parasitemia diluted to 1% every 3^rd^ day. Parasites cultured in DMEM supplemented with either vitamins or inorganic salts failed to support parasite growth and the parasitemia levels were similar to those observed in the cultures grown in DMEM alone after five cycles of propagation (Fig 1C). Supplementation of DMEM with missing amino acids was also insufficient to support the optimal growth of *B. duncani*, although the parasitemia levels at the end of cycle 5 were slightly higher in comparison to the parasitemia levels in DMEM (Fig 1C). The parasites grown in DMEM supplemented with either putrescine (+PUT) or lipids (LNA+LA) showed significantly (p <0.0001) higher parasitemia levels in comparison to the DMEM control (Fig 1C). Interestingly, at the end of cycle 5, the parasites grown in DMEM supplemented with either the combination of putrescine, lipids (LNA+LA) and six amino acids, or the combination of putrescine and lipids (LNA+LA) showed parasitemia levels similar to those observed in DMEM/F12, indicating that the combination of putrescine and lipids (LNA and LA) is sufficient to support optimal in vitro growth of *B. duncani* (Fig 1C). Similar analyses were conducted using drop-out media, which also identified putrescine, LNA and LA as key factors for parasite survival (Fig 1D).

**Fig 1.**
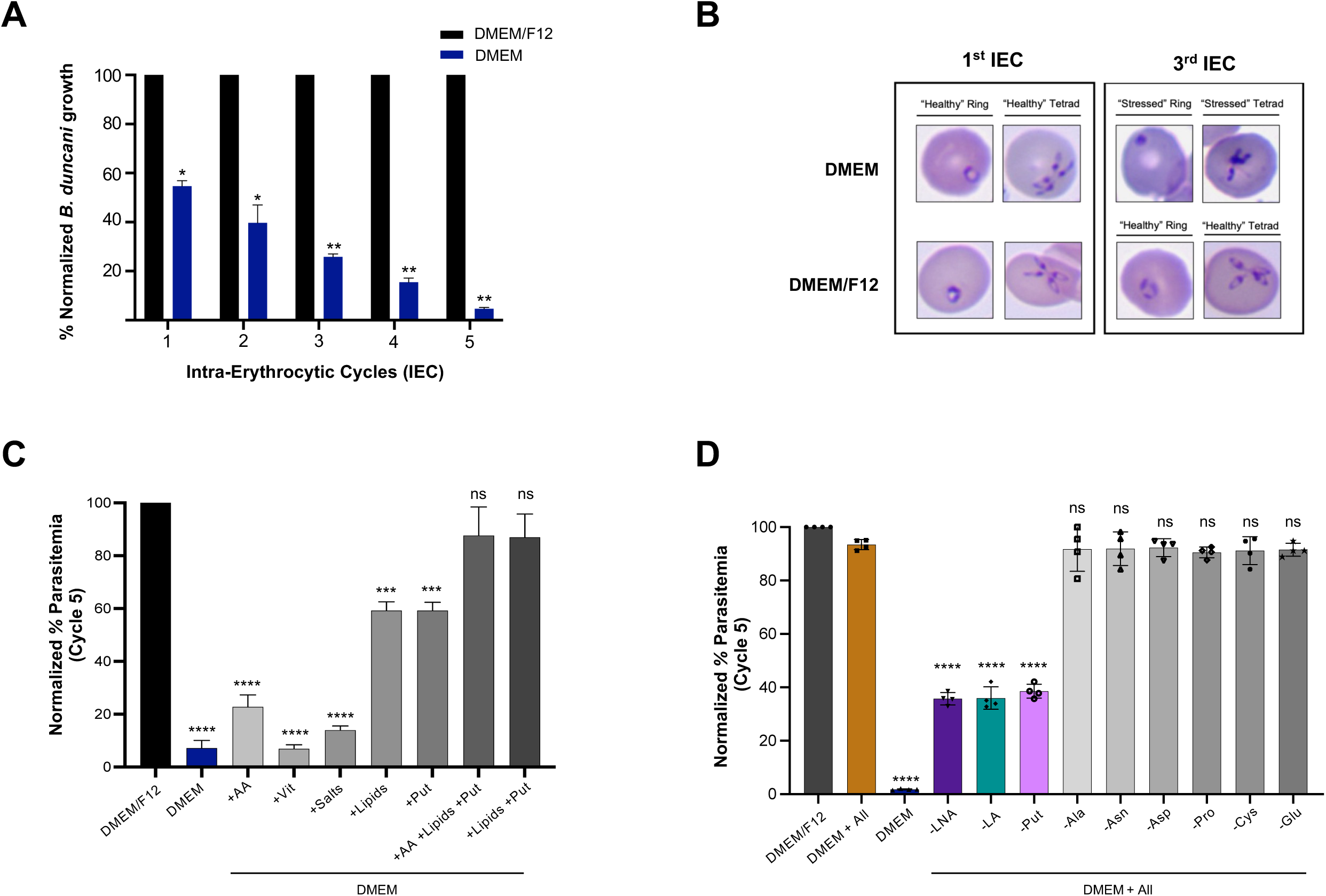
Essential nutrients for *B. duncani* in vitro growth in human RBCs. **A.** Comparison of *B. duncani* growth in DMEM/F12 and DMEM media over five 3-day propagation cycles. Parasitemia was monitored at the end of each 3-day cycle. A total of 3000-3500 RBCs were counted. The parasite growth in DMEM/F12 medium was used for normalization. Data presented as mean ± SD of two independent experiments performed in biological triplicates. Statistical significance of differences between parasite growth in DMEM/F12 and DMEM was estimated by multiple t-tests. * indicates significant P values <0.5, and ** indicates significant P values <0.01. **B.** Representative images of Giemsa-stained smears of *B. duncani* infected human RBCs at intraerythrocytic (IE) cycles 1 and 3 in DMEM or DMEM/F12 media showing healthy versus stressed rings and tetrads. **C.** Comparison of *B. duncani* growth in DMEM/F12, DMEM and DMEM supplemented with missing components including amino acids (AA), vitamins, salts, lipids (lipoic acid and linoleic acid), putrescine, combination of AA +Lipids +Put and combination of Lipids +Put at the end of cycle 5. The parasitemia was monitored at the end of each growth cycle. A total of 3000-3500 RBCs were counted. The parasite growth in DMEM/F12 medium was used for normalization. The normalized percent parasitemia at the end of cycle 5 is depicted in the graph. Data presented as mean ± SD of two independent experiments performed in biological triplicates. Statistical significance of differences between parasite growth in DMEM/F12 and other media was estimated by Welch’s t-test. * indicates significant P values <0.5, ** indicates significant P values <0.01, *** indicates significant P values <0.001, **** indicates significant P values <0.0001 and ns indicates no significant difference. **D.** Comparison of *B. duncani* growth in DMEM/F12, DMEM, DMEM containing all the missing components (DMEM + All) or DMEM+All depleted in individual components (linoleic acid (LNA), lipoic acid (LA), putrescine (Put), alanine ^12^, asparagine (Asn), aspartic acid (Asp), proline (Pro), cysteine (Cys) and glutamic acid (Glu)) over 5 cycles. The parasitemia was monitored at the end of each growth cycle. A total of 3000-3500 RBCs were counted. The parasite growth in DMEM/F12 medium was used for normalization. The normalized percent parasitemia at the end of cycle 5 is depicted in the graph. Data presented as mean ± SD of two independent experiments performed in biological triplicates. Statistical significance of differences between parasite growth in DMEM/F12 and other media was estimated by Welch’s t-test. * indicates significant P values <0.5, ** indicates significant P values <0.01, *** indicates significant P values <0.001, **** indicates significant P values <0.0001 and ns indicates no significant difference.

**Table I.** Differential components in DMEM, RPMI and DMEM/F12 media

|  |  | **DMEM**<br>Conc (mg/L) | **RPMI**<br>Conc (mg/L) | **DMEM/F12**<br>Conc (mg/L) |
| --- | --- | --- | --- | --- |
| **Amino Acids** | L-Alanine | - | - | 4.45 |
|  | L-Asparagine-H<sub>2</sub>O | - | 50 | 7.5 |
|  | L-Aspartic acid | - | 20 | 6.65 |
|  | L-Cysteine hydrochloride-H<sub>2</sub>O | - | - | 17.56 |
|  | L-Glutamic Acid | - | 20 | 7.35 |
|  | L-Proline | - | 20 | 17.27 |
| **Vitamins** | Biotin | - | 0.2 | 0.004 |
|  | Vitamin B12 | - | 0.005 | 0.68 |
| **Inorganic Salts** | Cupric sulfate (CuSO<sub>4</sub>.5H<sub>2</sub>O) | - | - | 0.0012 |
|  | Ferric sulfate (FeSO<sub>4</sub>.7H<sub>2</sub>O) | - | - | 0.42 |
|  | Magnesium Chloride (MgCl<sub>2</sub>) anhydrous | - | - | 28.57 |
|  | Zinc sulfate (ZnSO<sub>4</sub>.7H<sub>2</sub>O) | - | - | 0.43 |
| **Lipids** | Linoleic Acid | - | - | 0.044 |
|  | Lipoic Acid | - | - | 0.103 |
| **Other components** | Putrescine 2HCl | - | - | 0.081 |
|  | Hypoxanthine | - | - | 0.0112 |
|  | Thymidine | - | - | 0.001486 |

### *B. duncani* lacks the pathway for de novo polyamine biosynthesis and uses the salvage pathway for intraerythrocytic development and survival

The finding that putrescine is essential for intraerythrocytic development and survival of *B. duncani* led us to investigate the function of the machinery modulated by this polyamine and the molecular mechanism underlying its essential role. In other organisms, putrescine can either be synthesized de novo from arginine or transported from the cell environment via specialized polyamine transporters ^14^ (Fig 2A). De novo synthesis from arginine takes place along one of two enzymatically-driven biosynthetic routes, one relying on ornithine decarboxylase (ODC) and a second relying on arginine decarboxylase (ADC). To investigate whether a de novo polyamine biosynthesis pathway exists in *B. duncani* WA-1, we examined the susceptibility of the parasite to DL-α-difluoromethylornithine (DFMO), an inhibitor of ODC enzyme ^15^, and DL-alpha-difluoromethylarginine (DFMA), an inhibitor of ADC enzyme ^16^, in media lacking or supplemented with putrescine. Both drugs failed to inhibit the growth of *B. duncani* in both media (Fig. 2B and C). Consistent with these pharmacological data, supplementation with L-Ornithine up to 1 mg/ml did not rescue the growth defect resulting from putrescine depletion (Figure 2D). Together these results suggest that *B. duncani* does not biosynthesize putrescine de novo, and therefore must rely solely on the putrescine salvage pathway for survival within human erythrocytes.

**Fig 2.**
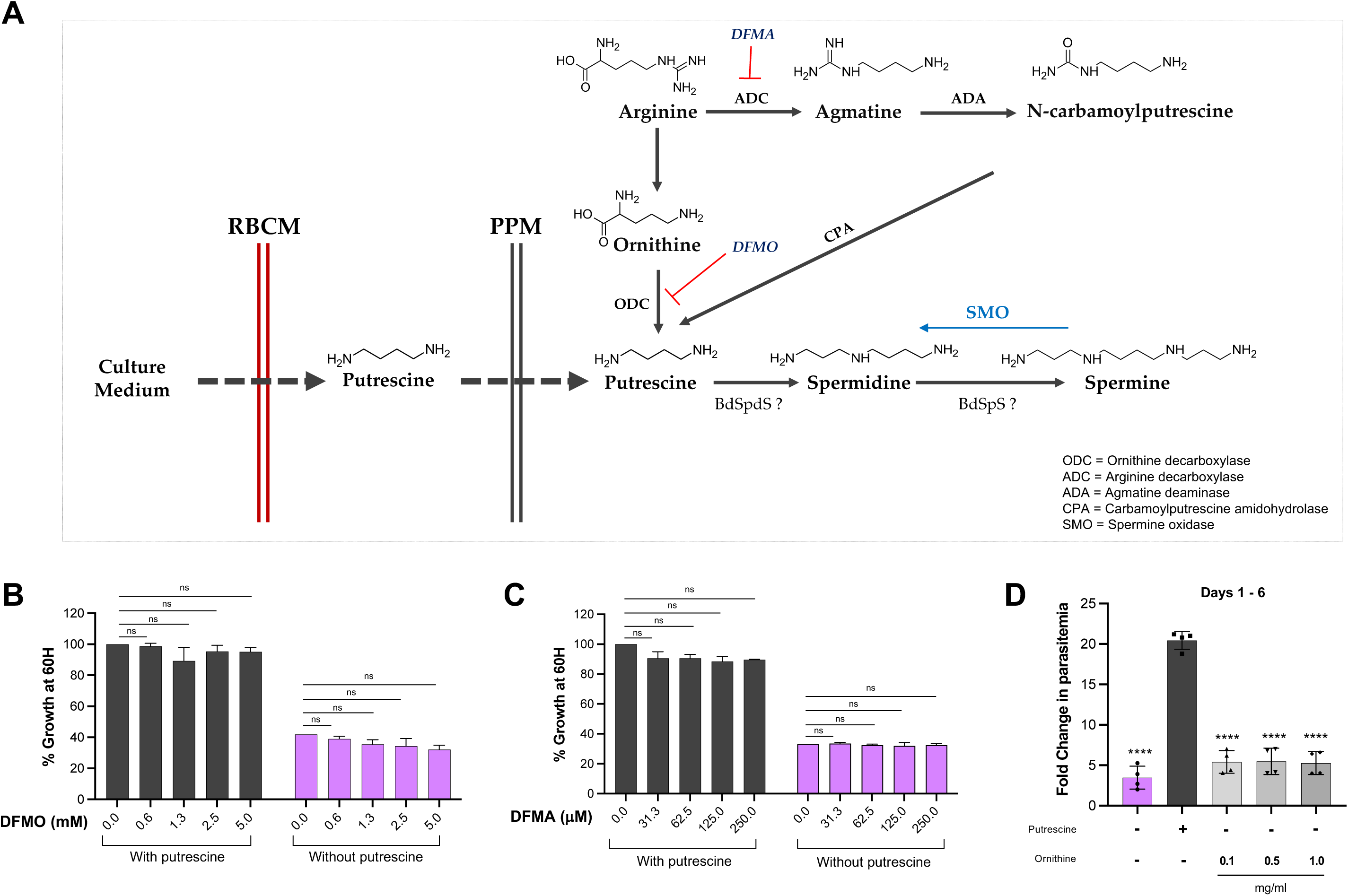
Polyamine salvage pathway is critical for optimal in vitro growth of *B. duncani*. **A.** Schematic representation of de novo biosynthesis and salvage pathways of polyamine metabolism. **B.** In vitro efficacy of DL-α-difluoromethylornithine (DFMO) on *B. duncani* grown in the presence or absence of putrescine. Data presented as mean ± SD of three independent experiments performed in biological triplicates. Statistical significance of differences was estimated by Welch’s t-test. ns indicates no significant difference. **C.** In vitro efficacy of DL-α-difluoromethylarginine (DFMA) on *B. duncani* grown in the presence or absence of putrescine. Data presented as mean ± SD of three independent experiments performed in biological triplicates. Statistical significance of differences was estimated by Welch’s t-test. ns indicates no significant difference. **D.** Ornithine supplementation fails to restore growth defect in parasites cultured in putrescine depleted medium. *B. duncani* in vitro growth in putrescine containing medium, putrescine depleted medium, or putrescine depleted medium supplemented with 0.1 mg/ml, 0.5 mg/ml and 1 mg/ml ornithine was monitored for a 6-day period. The parasitemia was monitored on day 1 and day 6. Fold change in parasitemia was calculated by counting a total of 3000-3500 RBCs. Data presented as mean ± SD of two independent experiments performed in biological triplicates. Statistical significance of differences between parasite growth in putrescine containing or lacking medium and putrescine depleted media supplemented with different concentrations of ornithine were estimated by Welch’s t-test. **** indicates significant P values <0.0001.

### Putrescine depletion in *B. duncani* results in elevated levels of reactive oxygen species

Elevated intracellular reactive oxygen species (ROS) levels indicates oxidative stress and often presages stunted growth and associated death ^17, 18^. To determine whether parasites grown in the absence of putrescine were indeed under oxidative stress, *B. duncani*-infected RBCs were stained with dihydrorhodamine 123 (DHR123), and Hoechst 33342 (a nuclear stain), and the resulting fluorescence was compared to that of parasites grown in the presence of putrescine, spermidine or spermine. DHR123 is an uncharged non-fluorescent molecule that is cleaved inside the cells into its fluorescent byproduct rhodamine via the action of ROS ^19^. Rhodamine-positive *B. duncani*-infected RBCs in cultures maintained in media lacking or supplemented with polyamines were examined by fluorescence microscopy (Fig. 3A) and quantified using flow cytometry (Fig. 3B). *B. duncani* cultures maintained in the absence of putrescine showed 5x, 10x and 20x more rhodamine-positive *B. duncani*-infected RBCs compared to those maintained in the presence of putrescine, spermidine or spermine, respectively (Fig 3B). As a control, the antiparasitic drug, artemisinin, which has been shown to exert its cytotoxic effect via production of free radicals and ROS in other apicomplexan parasites ^20^, induced a dose-dependent increase in ROS production in *B. duncani*-infected RBCs even in the presence of putrescine (Fig. 3B). Combined fluorescence and brightfield microscopic examination of parasites maintained in the above-mentioned growth media also displayed similar patterns, with the majority of parasites maintained in putrescine, spermidine, and spermine-supplemented medium staining negative for rhodamine, whereas the majority of parasites maintained in putrescine-depleted medium stained positive for rhodamine (Fig. 3A).

**Fig 3.**
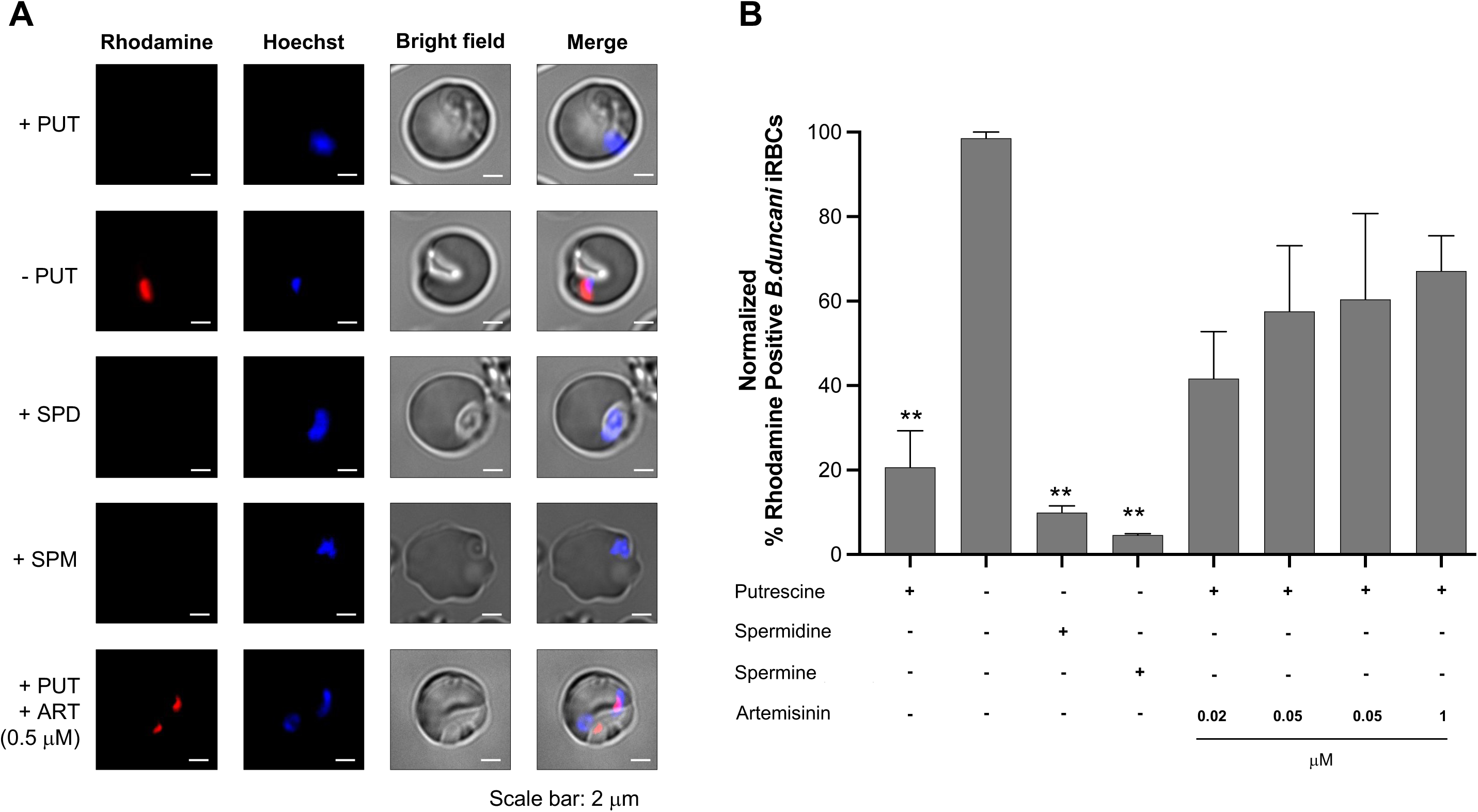
Reactive oxygen species (ROS) levels are high in *B. duncani* grown in putrescine depleted medium. **A.** Representative live cell fluorescence images showing rhodamine signal (red) post incubation with dihydrorhodamine 123 (DHR123), an uncharged non-fluorescent molecule that is cleaved into its fluorescent byproduct rhodamine via the action of ROS inside the cells) in *B. duncani* cultured in different growth media, including putrescine depleted medium (-PUT) or putrescine depleted media supplemented with putrescine (1mg/ml; 11.3 mM) (+PUT), spermidine (1mg/ml; 6.9 mM) (+SPD) and spermine (1mg/ml; 4.9 mM) (+SPM), or putrescine supplemented (1mg/ml; 11.3 mM) medium containing 0.5 μM of artemisinin (+ PUT + ART). Hoechst (blue) is used to stain the parasite nuclei. Scale bar 2μm. Data presented as mean ± SD of three independent experiments performed in biological triplicates. **B.** Comparison of the percent of rhodamine positive parasites in different growth media. The parasites cultured in different media were incubated with dihydrorhodamine123 for 15 minutes and subsequently rhodamine signal was measured using flow cytometry after washing. Percent rhodamine positive parasites in putrescine depleted medium was used for normalization. Data presented as mean ± SD of three independent experiments performed in biological triplicates. Statistical significance of differences between percent rhodamine positive parasites in putrescine depleted medium and putrescine depleted media supplemented with putrescine, spermidine or spermine were estimated by Welch’s t-test. ** indicates significant P values <0.01.

### Spermidine is the key polyamine essential for optimal growth of *B. duncani*

Our finding that putrescine uptake is essential for *B. duncani* intraerythrocytic development and survival led us to investigate whether putrescine itself or one or both of its derivatives spermidine and spermine are the key polyamines required for parasite viability (Fig. 4A). We first examined whether the growth defect caused by putrescine depletion could be rescued by the addition of spermidine or spermine. As shown in Fig. 4B, parasites propagated in media lacking putrescine but supplemented with either spermidine or spermine were viable and replicated at similar rates as those grown in the presence of putrescine (Fig. 4B). We then assessed the effect of inhibition of the parasite spermidine synthase (SPDS) by 4-methylcyclohexylamine (4-MCHA) and spermine oxidase (SMO) by MDL 72527 on parasites grown in media either lacking putrescine or putrescine depleted media supplemented with either putrescine, spermidine or spermine. The half-maximal inhibitory concentration (IC_50_) values of 4-MCHA in putrescine depleted or containing media were found to be similar (Fig. 4C and table II). However, a ∼7.5 fold increase in the IC_50_ of 4-MCHA was calculated for parasites grown in putrescine depleted media but supplemented with either spermidine or spermine (Fig 4C and table II). On the other hand, the IC_50_ values of MDL 72527 for parasites grown in putrescine depleted or containing media as well as spermine-supplemented medium were found to be similar (Fig. 4D). Interestingly, a ∼3.5-fold increase in MDL 72537 IC_50_ value was observed in parasites cultured in spermidine-supplemented medium (Fig. 4D and table III). A major shift was observed in the IC_50_ values of 4-MCHA and MDL 72527 specifically in parasite cultures maintained in putrescine-depleted medium but supplemented with spermidine. Altogether, these findings conclusively indicate that spermidine is the key polyamine required for parasite growth.

**Fig 4.**
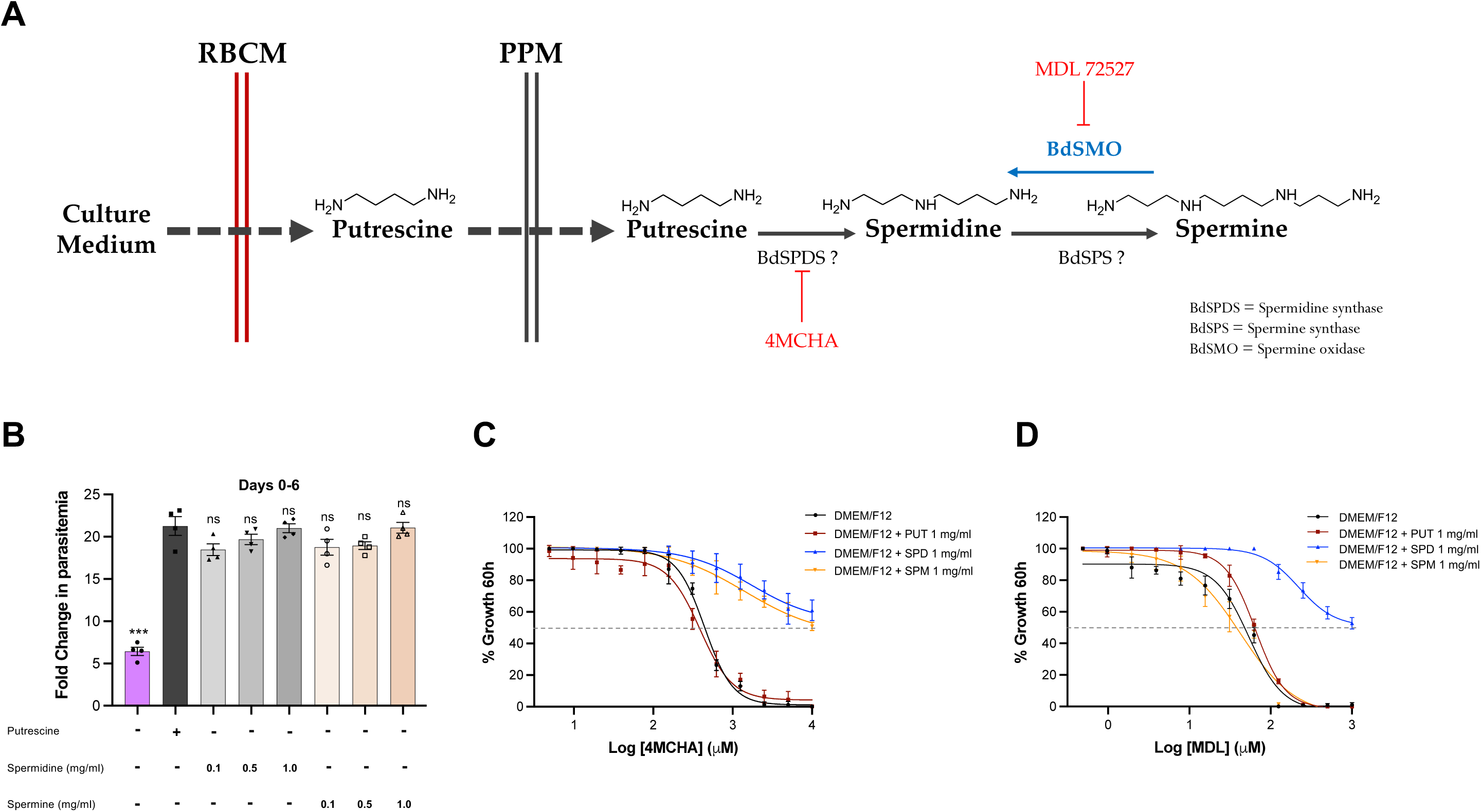
Spermidine is the key polyamine required for intraerythrocytic growth of *B. duncani* in human RBCs. **A.** Schematic representation of polyamine salvage pathway in *B. duncani*. **B.** Spermidine and spermine supplementation restores growth defect in parasites grown in putrescine depleted medium. *B. duncani* in vitro growth in putrescine containing medium, putrescine depleted medium, or putrescine depleted medium supplemented with 0.1 mg/ml, 0.5 mg/ml, and 1 mg/ml of either spermidine or spermine was monitored for a 6-day period. Fold change in parasitemia was calculated by counting a total of 3000-3500 RBCs on the 6^th^ day. Data presented as mean ± SD of two independent experiments performed in biological triplicates. Statistical significance of differences between parasite growth in putrescine containing or lacking medium and putrescine depleted medium supplemented with different concentrations of spermidine and spermine were estimated by Welch’s t-test. *** indicates significant P values <0.001 and ns indicates no significant difference. **C.** In vitro efficacy and IC_50_ determination of trans-4-methylcyclohexylamine (4-MCHA); a spermidine synthase inhibitor, against *B. duncani* in DMEM/F12 and DMEM/F12 supplemented with 1 mg/ml of either putrescine or spermidine or spermine. Data presented as mean ± SD of three independent experiments performed in biological triplicates. The IC_50_ values of 4-MCHA under growth in different media are displayed in the table below the graph. **D.** In vitro efficacy and IC_50_ determination of MDL 72527; a spermine oxidase inhibitor, against *B. duncani* in DMEM/F12 and DMEM/F12 supplemented with 1 mg/ml of either putrescine or spermidine or spermine. Data presented as mean ± SD of three independent experiments performed in biological triplicates. The IC_50_ values of MDL 72527 under growth in different media are displayed in the table below the graph.

**Table II.** IC_50_ of 4-MCHA in B. duncani in different growth media

| Growth Medium | 4-MCHA IC<sub>50</sub> (μM) |
| --- | --- |
| DMEM/F12 | 444.1 |
| DMEM/F12 + PUT 1 mg/ml | 388.4 |
| DMEM/F12 + SPD 1 mg/ml | 3127 |
| DMEM/F12 + SPM 1 mg/ml | 2872 |

**Table III.** IC_50_ of MDL 72527 in *B. duncani* in different growth media

| Growth Medium | MDL 72527 IC<sub>50</sub> (μM) |
| --- | --- |
| DMEM/F12 | 54.26 |
| DMEM/F12 + PUT 1 mg/ml | 64.70 |
| DMEM/F12 + SPD 1 mg/ml | 221.2 |
| DMEM/F12 + SPM 1 mg/ml | 50.41 |

### Polyamine-mediated transcriptional regulation in *B. duncani*

To investigate the impact of putrescine depletion on gene expression in *B. duncani*, the parasites were cultured in media lacking or supplemented with putrescine for 24 hours and their transcription profiles analyzed by Illumina RNA-seq. The normalized counts and rlog-transformed data were generated using DESeq2 analysis ^21, 22^ and a heatmap was constructed (Fig. 5A). Out of 4,222 genes encoded by *B. duncani*, 66 genes were found to be differentially expressed between parasites cultured in the absence or presence of putrescine (Fig. 5A and 5B) with 37 genes found to be downregulated and 29 genes upregulated in parasites propagated in the absence of putrescine compared to those maintained in the presence of putrescine (Fig. 5B). Gene ontology biological process analysis of these differentially expressed genes identified several different biological processes affected by the putrescine supplementation or depletion. Half of the downregulated genes in parasites cultured in the absence of putrescine were found to encode hypothetical proteins (Fig. 5C). Interestingly, of the downregulated genes with predicted functions 17% were found to play role in protein translation, 6% in protein phosphorylation and 3% in either ATP binding, DNA binding, phosphoinositide pathway, regulation of mRNA stability, nuclear-transcribed mRNA catabolism, regulation of DNA regulation, hydrolase activity and inositol phosphate biosynthesis (Fig. 5C). Out of the genes upregulated under putrescine depletion, 32% encode hypothetical proteins, 32% are involved in nuclear-transcribed mRNA catabolism, and 6% involved in protein folding, intracellular protein transport, carbohydrate derivate metabolism, mRNA splicing via spliceosomes, negative regulation of translation and cellular glucose metabolism (Fig. 5D).

**Fig 5.**
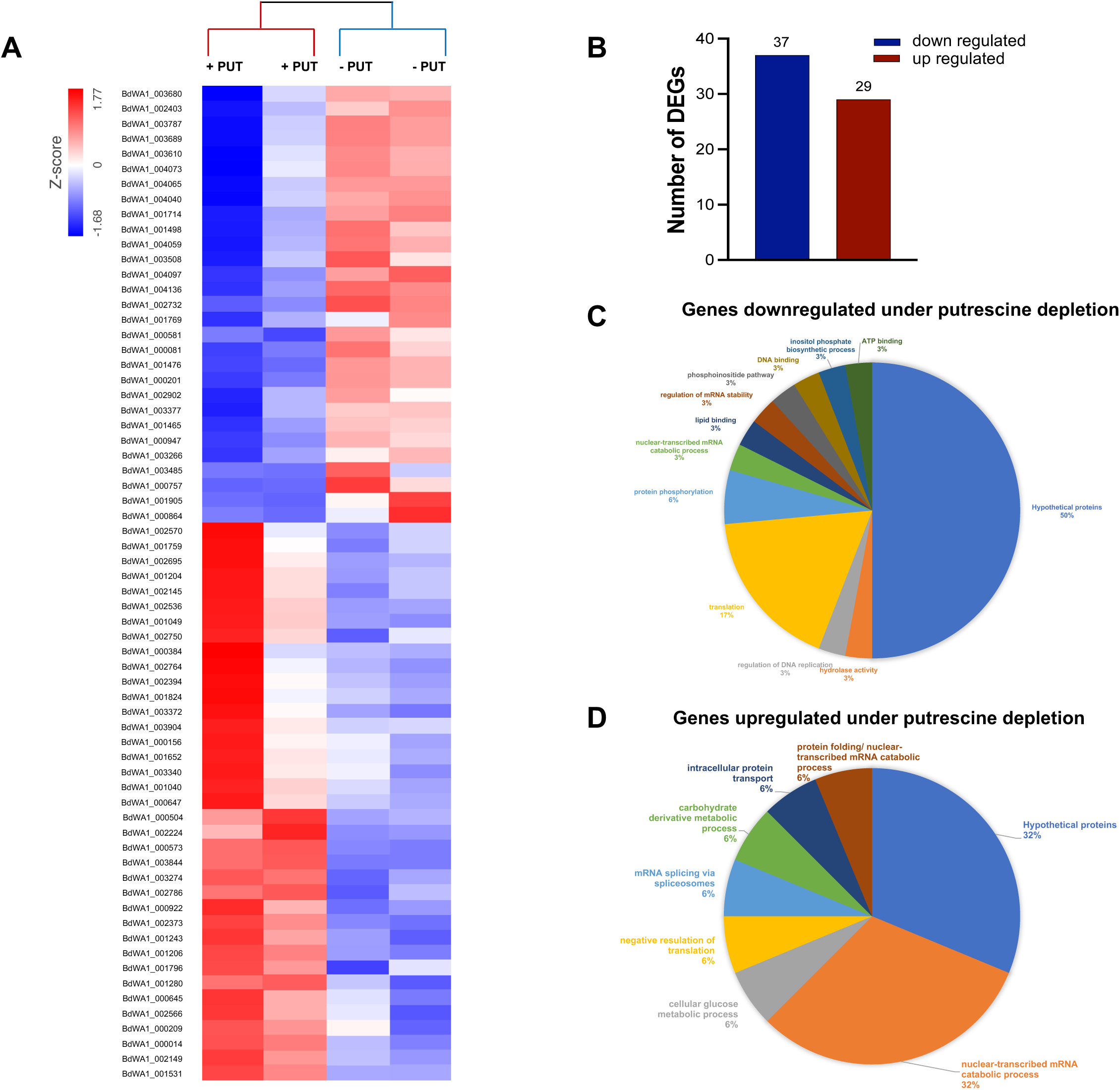
RNA-seq expression data analysis in *B. duncani* parasites cultured in the presence or absence of putrescine for 24 H display differential expression of genes. **A.** Heatmap and hierarchical clustering of log 2-fold changes in gene expression in *B. duncani* in response to putrescine depletion. Data is presented for two independent biological replicates. **B.** *B. duncani* differentially expressed genes (DEGs) from parasites propagated in the absence or presence of putrescine. The upregulated and downregulated genes are colored blue and maroon, respectively. **C.** Gene ontology pathways associated with the genes downregulated upon putrescine depletion. **D.** GO pathways associated with the genes upregulated upon putrescine depletion.

### Spermidine-mediated hypusination of the elongation factor eIF5A is critical for *B. duncani* intraerythrocytic growth

Studies of other organisms have revealed that spermidine acts as a donor of the amino-butyl group to lysine 51 of the elongation factor 5A (eIF5A) to form hypusine (N^ε^-[4-amino-2-hydroxybutyl]-lysine) ^23^. This post-translational hypusine synthesis is a two-step enzymatic process, catalyzed by deoxyhypusine synthase (DHS) and deoxyhypusine hydrolase (DOHH). To investigate the importance of *B. duncani* DHS enzyme in parasite development, we examined the susceptibility of the parasite to N1-guanyl-1,7-diaminoheptane (GC7), a spermidine analog that inhibits DHS and prevents eIF5A hypusination (Fig. 6A). The compound was found to be effective against the parasite in putrescine containing medium (DMEM/F12) with an IC_50_ of ∼120 µM (Fig. 6B). Supplementation of the medium with putrescine, spermidine or spermine had no effect on the potency of the drug (Fig. 6B). We next examined the effect of GC7 treatment on *B. duncani* BdeIF5A hypusination using anti-eIF5A and anti-hypusine antibodies on parasite lysates from *B. duncani* cultures grown in the presence or absence of GC7. Whereas the levels of eIF5A remained unchanged in the presence or absence of GC7 (Fig. 6C, 6D), the levels of hypusinated eIF5A in the presence of GC7 were ∼20% in comparison to those found in the absence of the compound (Fig. 6C, 6E). As a control, the levels of BdHsp70-2 were unchanged in the absence or presence of GC7. Together, these results indicate that a hypusination pathway is present in *B. duncani* and its inhibition blunts the hypusination of eIF5A. To assess whether the growth defect observed upon putrescine depletion in *B. duncani* is due to altered eIF5A hypusination, immunoblot analyses were conducted using anti-eIF5A and anti-hypusine antibodies on parasite lysates from cultures grown in media lacking putrescine or supplemented with either putrescine, spermidine or spermine (Fig. 6F-H). An approximate 90% reduction in the levels of hyp-eIF5A was measured in parasites cultured in the absence of putrescine compared to parasites cultured in the presence of this polyamine, whereas no significant differences were measured in the levels of eIF5A (Fig. 6F and 6G). Supplementation with spermidine or spermine restored Hyp-eIF5A levels similar to those measured in the presence of putrescine (Fig. 6F and 6G). Together, these studies demonstrate that spermidine-mediated hypusine formation is impacted when the parasites are cultured in putrescine lacking growth medium. Specifically, the failure to form a hypusine residue on lysine 51 of eIF5A impacts protein translation and thus parasite growth.

**Fig 6.**
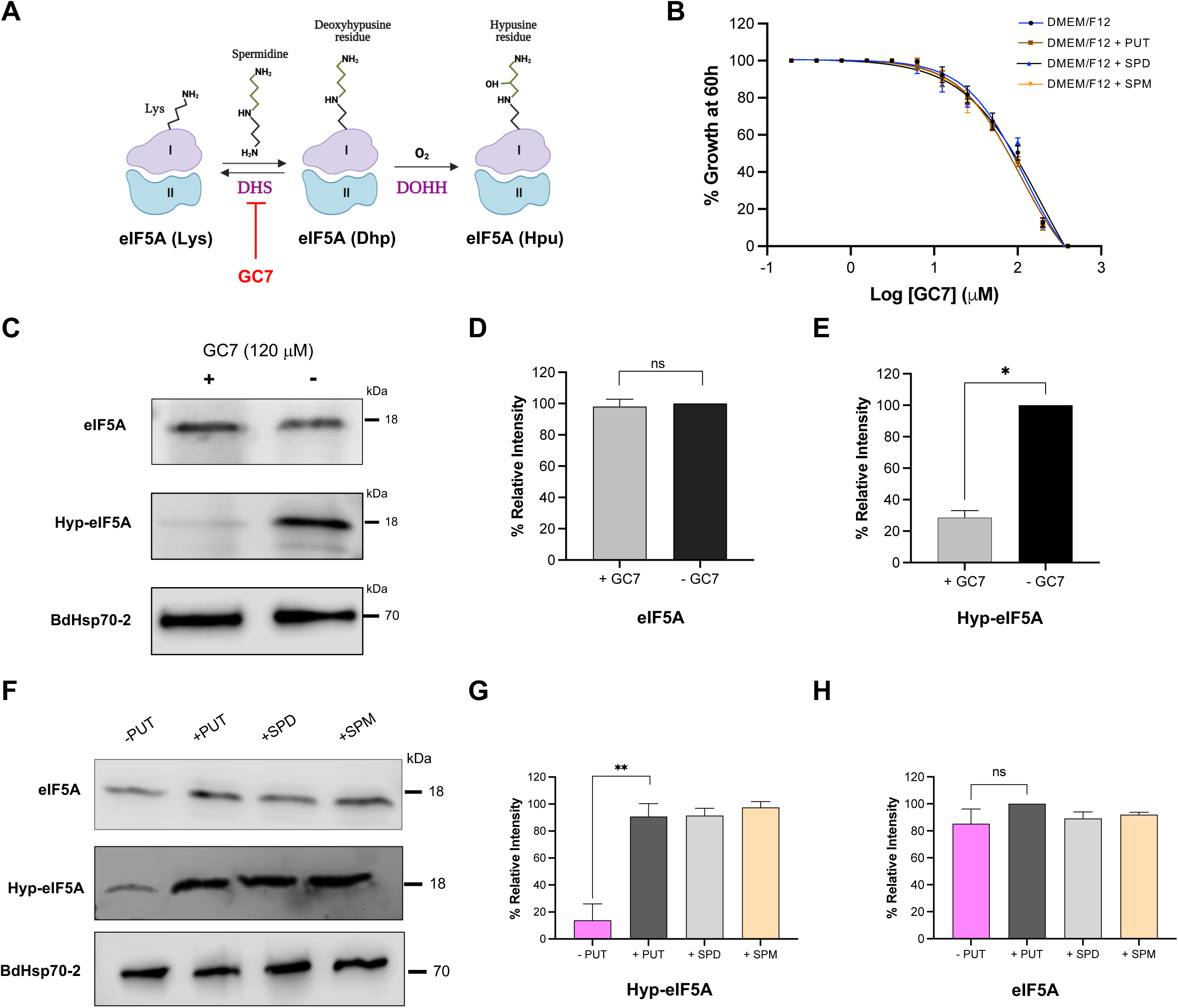
*B. duncani* parasites grown in putrescine depleted medium display reduced levels of eIF5A hypusination. **A.** Schematic representation of the eIF5A hypusination pathway. **B.** In vitro efficacy and IC_50_ determination of GC7; a deoxyhypusine synthase (DHS) inhibitor, against *B. duncani* in DMEM/F12 and DMEM/F12 supplemented with 1 mg/ml of either putrescine or spermidine or spermine. Data presented as mean ± SD of three independent experiments performed in biological triplicates. **C.** Western blot analysis using anti-eIF5A and anti-hypusine Abs against lysates from parasites grown in the presence or absence of GC7 IC_50_ concentration (120 μM) for 48 H. Anti-Bdhsp70-2 antiserum was used as loading control. **D.** The graph showing the relative levels of eIF5A in parasite lysates from untreated or GC7 treated parasites. Data presented as mean ± SD of three independent experiments. **E.** The graph showing the relative levels of hypusinated-eIF5A (Hyp-eIF5A) in parasite lysates from untreated or GC7 treated parasites. Hypusination of eIF5A (Hyp-eIF5A) is ∼80% less in parasites grown in the presence of GC7 inhibitor in comparison to the control untreated parasites. Data presented as mean ± SD of three independent experiments. **F.** Western blot analysis using anti-eIF5A and anti-hypusine Abs against lysates from parasites cultured in putrescine depleted (-PUT) or containing medium (+PUT), and putrescine depleted medium supplemented with either spermidine (+SPD) or spermine (+SPM) for 48 H. Anti-Bdhsp70-2 antiserum was used as loading control. **G.** The graph showing the relative levels of hypusinated-eIF5A (Hyp-eIF5A) in parasite lysates from untreated or GC7 treated parasites. Hypusination of eIF5A (Hyp-eIF5A) is ∼90% less in parasites grown in -PUT medium in comparison to the parasites grown in +PUT medium. Data presented as mean ± SD of three independent experiments. **H.** The graph showing the relative levels of eIF5A in parasite lysates from untreated or GC7 treated parasites. Data presented as mean ± SD of three independent experiments. Statistical significance of differences were calculated using Welch’s t-test. * indicates significant P values <0.05, ** indicates significant P values <0.001 and ns indicates no significant difference.

### Spermidine-mediated hypusination is a conserved mechanism that is essential for intraerythrocytic development of apicomplexan parasites

Several apicomplexan parasites including the human malaria parasite *P. falciparum* can grow in the absence of exogenous polyamines and possess a de novo polyamine biosynthesis pathway (Fig. 7A). The intraerythrocytic stages of *P. falciparum* 3D7 are sensitive to inhibitors of the polyamine biosynthesis pathway, including DFMO (targets ODC) and 4-MCHA (targets SPDS), as well as to the inhibitor of eIF5A hypusination pathway, GC7 (targets DHS) (Fig. S2). The eIF5A hypusination pathway is well conserved in *P. falciparum* (Fig. 7B). The inhibition of ODC by DFMO can be reversed by the addition of putrescine, spermidine and spermine (Fig. S2 A). On the other hand, the inhibition of SPDS by 4-MCHA could be reversed by the addition of spermidine or spermine, but not putrescine (Fig. S2 B). To assess whether blocking de novo putrescine biosynthesis pathway through the inhibition of ODC in *P. falciparum* results in defects in the hypusination of eIF5A, immunoblot analyses were performed on parasite lysates prepared from cultures grown in the absence or presence of DFMO (IC_50_ concentration), or in the presence of DFMO but supplemented with putrescine, spermidine or spermine, using anti-eIF5A and anti-hypusine antibodies. No differences were observed in the levels of eIF5A in parasites cultured in the aforementioned growth media (Fig. 7C and 7D). However, a ∼90% reduction in the levels of hyp-eIF5A was observed in the parasites treated with DFMO (IC_50_ concentration) in comparison to the untreated parasites (Fig. 7C and 7E). Interestingly, DFMO-treated parasites cultured in presence of putrescine, spermidine and spermine displayed Hyp-eIF5A levels similar to those of untreated parasites (Fig. 7C and 7E). GAPDH was used as loading control (Fig. 7C). Furthermore, the effect of DHS inhibition by GC7 treatment on eIF5A hypusination in *P. falciparum* was tested. The *P. falciparum* parasites grown in the absence or presence of GC7 (IC_50_ concentration) displayed similar levels of eIF5A using Western blotting (Fig. 7F, 7G). GC7 treated parasites cultured in the presence of putrescine, spermidine and spermine also displayed levels of eIF5A similar to those of the GC7 treated parasites grown in the absence of these polyamines (Fig. 7F, 7G). The levels of Hyp-eIF5A levels in GC7 treated *P. falciparum* parasites were ∼80% lower than in the untreated parasites (Fig. 7F and 7H). GC7 treated parasites cultured in the presence of putrescine, spermidine and spermine displayed Hyp-eIF5A levels similar to those of the untreated parasites (Fig. 7F and 7H), indicating that the GC7 mediated inhibition of DHS can be reversed through the addition of excess polyamines.

**Fig 7.**
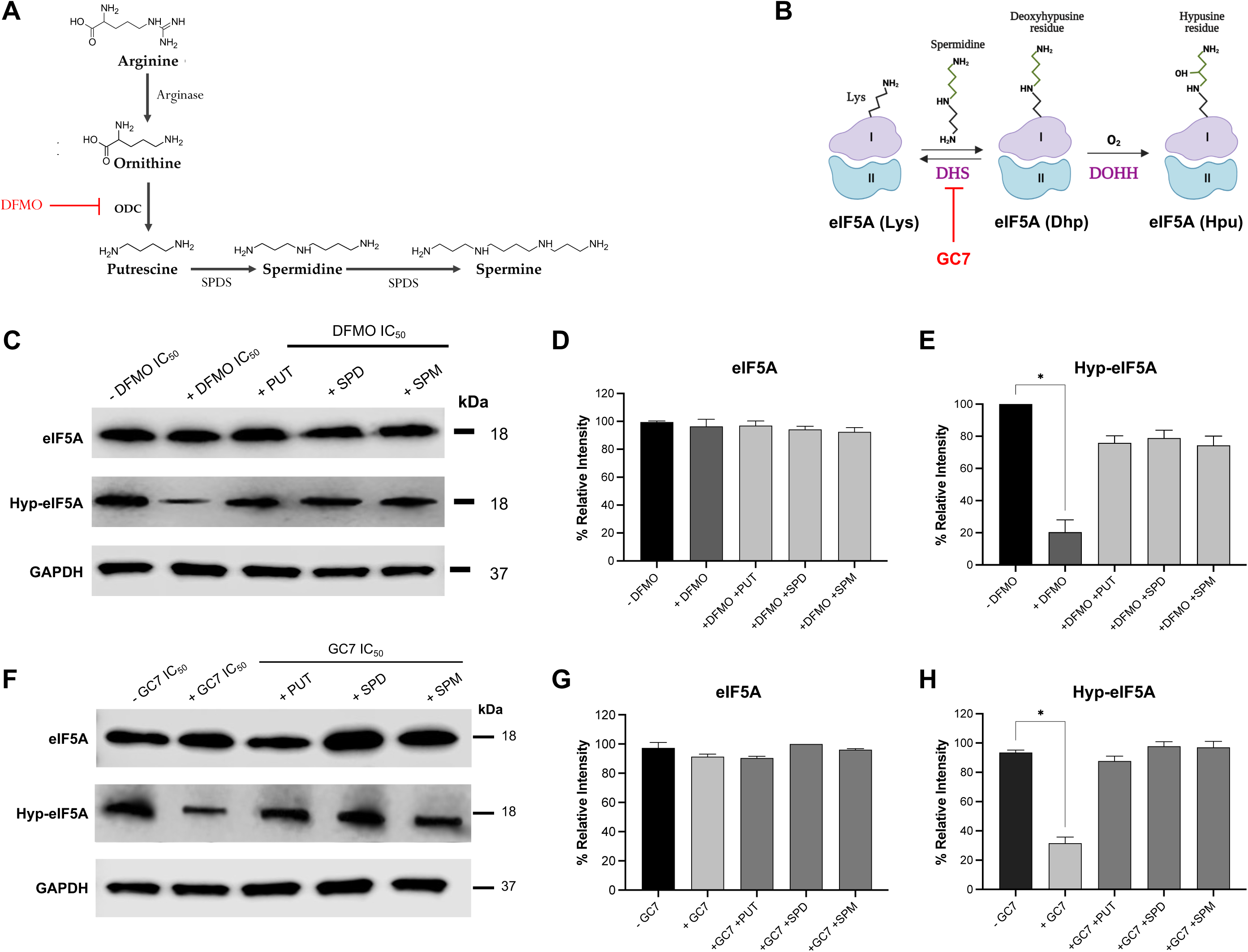
*P. falciparum* 3D7 parasites treated with DFMO and GC7 display reduced levels of eIF5A hypusination. **A.** Schematic representation of the de novo pathway of polyamine synthesis in *P. falciparum*. **B.** Schematic representation of the eIF5A hypusination pathway. **C.** Western blot analysis using anti-eIF5A and anti-hypusine Abs against lysates from parasites cultured in the absence or presence of DFMO (IC_50_ concentration, 1mM), or in the presence of DFMO (IC_50_ concentration) but supplemented with 1mg/ml of putrescine (+PUT), or 1mg/ml of spermidine (+ SPD), or 1mg/ml of spermine (SPM) for 72 H. Anti-GAPDH Ab was used as loading control. **D.** The graph depicting the relative levels of eIF5A in parasite lysates from different culture conditions. Data presented as mean ± SD of three independent experiments. **E.** The graph showing the relative levels of hypusinated-eIF5A (Hyp-eIF5A) in parasite lysates from different culture conditions. Data presented as mean ± SD of three independent experiments. Levels of hypusinated-eIF5A (Hyp-eIF5A) are ∼80% less in parasites grown in the presence of DFMO (IC_50_ concentration) in comparison to the control untreated parasites. **F.** Western blot analysis using anti-eIF5A and anti-hypusine Abs against lysates from parasites cultured in the absence or presence of GC7 (IC_50_ concentration; 15μM), or in the presence of GC7 (IC_50_ concentration) but supplemented with 1mg/ml of putrescine (+PUT), or 1mg/ml of spermidine (+ SPD), or 1mg/ml of spermine (SPM) for 72 H. Anti-GAPDH Ab was used as loading control. **G.** The graph depicting the relative levels of eIF5A in parasite lysates from different culture conditions. Data presented as mean ± SD of three independent experiments. **H.** The graph showing the relative levels of hypusinated-eIF5A (Hyp-eIF5A) in parasite lysates from different culture conditions. Data presented as mean ± SD of three independent experiments. Levels of hypusinated-eIF5A (Hyp-eIF5A) are ∼70% less in parasites grown in the presence of GC7 (IC_50_ concentration) in comparison to the control untreated parasites.

## Discussion

The polyamines putrescine, spermidine and spermine are organic polycationic amines that play essential roles in all eukaryotes including pathogenic protozoa ^24, 25^. While some parasites employ de novo synthesis routes for production of these polyamines; others rely on the uptake of preformed polyamines for survival. However, in most parasites, it remains unclear which of the three polyamines are critical for parasite development and by what mechanism. In this study, we provided the first evidence that *B. duncani* exclusively relies on a salvage pathway for polyamine biosynthesis and demonstrated that the key output of this pathway is spermidine, which plays a critical role in the operation of the translation machinery through hypusination of eIF5A (Fig. 8). The availability of full genome sequences from several protozoan parasites and in some cases further genetic and pharmacological studies have helped map the components of the polyamine biosynthesis pathway in *Plasmodium* spp., *Trypanosoma* spp., *Toxoplasma* spp. and *Cryptosporidium* spp. ^26^. In organisms primarily reliant on de novo biosynthesis is the main route for synthesis of polyamines, the pathway initiates with the conversion of arginine to ornithine by Arginase 1 (ARG1). Ornithine is further decarboxylated to form putrescine by the action of an ornithine decarboxylase (ODC), a rate-limiting step in the biosynthesis of polyamines. Putrescine is further converted first to spermidine and then to spermine by the actions of spermidine synthase (SPDS) and spermine synthase (SPS) respectively, through the sequential addition of aminopropyl groups (Fig. 8). In *T. cruzi* (the causative agent of chagas disease in humans), a functional ODC is absent, and putrescine is either taken up from the host or (under polyamine shortage conditions) is formed by an alternate route through the conversion of arginine into agmatine by an arginine decarboxylase (ADC) ^27, 28^. Agmatine then serves as a precursor for the synthesis of N-carbamoyl-putrescine, which is later converted into by putrescine by carbamoylputrescine amidohydrolase (CPA) (Fig. 2A). The loss of a functional ODC in certain parasites such as *T. cruzi* and *T. gondii* is a classic example of reductive genome evolution ^29^ where the parasites have lost the genes involved in de novo polyamine biosynthesis pathway and rely on their host for polyamine salvage. In this study, we showed that the intraerythrocytic stages of *B. duncani* parasites are insensitive to inhibition by the ODC inhibitor DFMO. Furthermore, we found that ornithine supplementation failed to rescue parasite growth on media lacking polyamines, indicating that a functional ODC is absent in *B. duncani*. Accordingly, bioinformatic analyses failed to identify an *ODC* gene or ODC protein in the genome or proteome of *B. duncani*, respectively. Together these observations suggest that *B. duncani* has undergone reductive evolution by losing de novo polyamine biosynthesis pathway and relies on its host to salvage polyamines.

**Fig 8.**
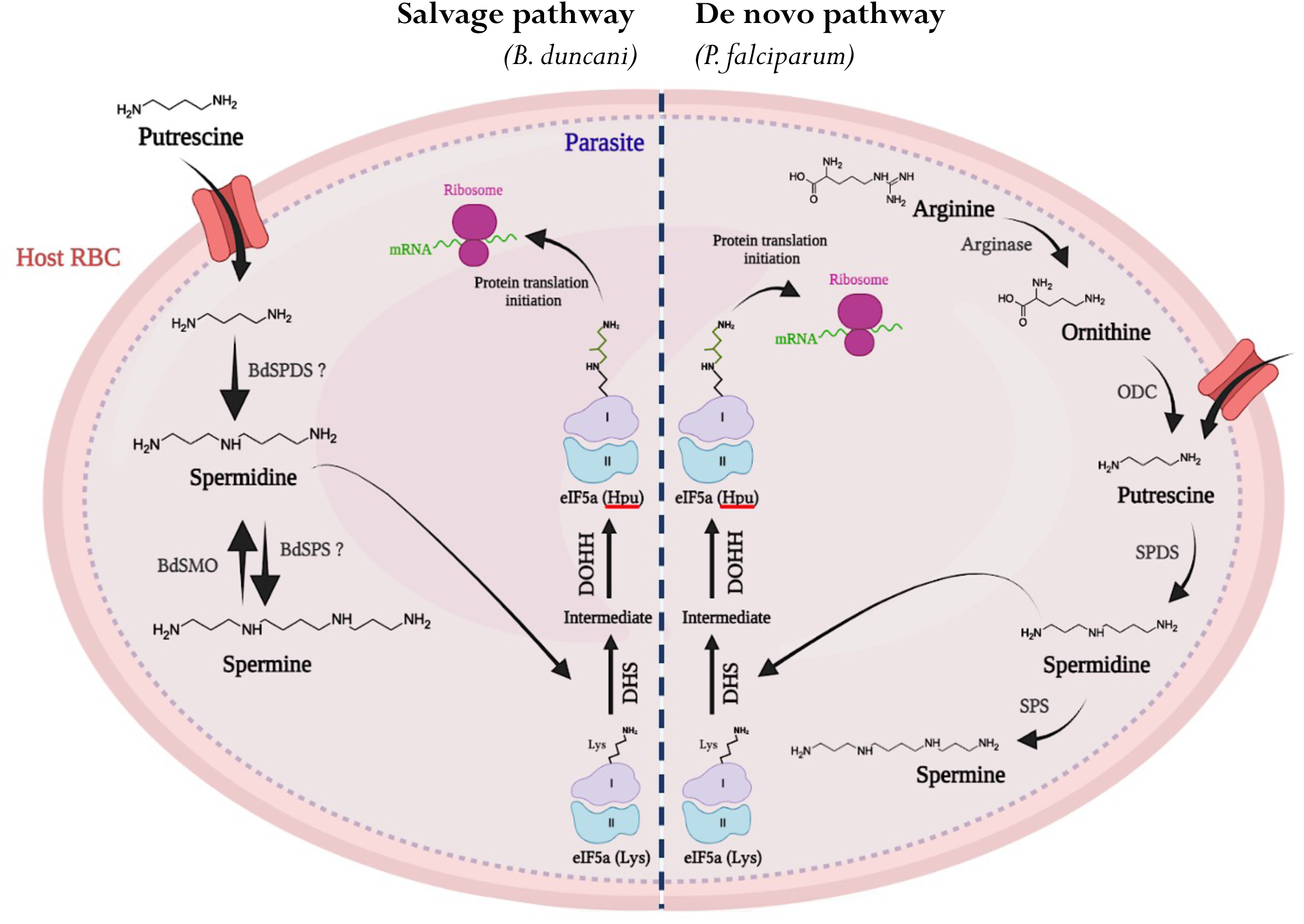
Schematic model showing the control of eIF5A hypusination through polyamine pathway. In *B. duncani*, putrescine is salvaged and utilized for the synthesis of spermidine and spermine. In *P. falciparum*, putrescine biosynthesis commences through the conversion of arginine into ornithine through the action of arginase. Ornithine is converted into putrescine through the action of ornithine decarboxylase (ODC). Putrescine is processed further to form spermine and spermine. In both *B. duncani* and *P. falciparum*, spermidine is used as a substrate in the hypusination pathway and results in the addition of hypusine residue on eIF5A. Hypusinated eIF5A is used for the initiation of protein translation.

Although, *ODC* is absent in *B. duncani*, the parasite can convert putrescine into spermidine and spermine, indicating the presence of spermidine synthase (*SPDS*) and spermine synthase (*SPS*) enzymatic activity. Although, canonical *SPDS* and *SPS* genes have not yet been identified in *B. duncani* genome, in silico analyses identified a protein BdWA1_000190 which shares 36% identity with a *Theileria parva* putative protein TpMuguga_01g02015 annotated as SPDS. Whether this enzyme catalyzes this dual activity remains to be determined. In some organisms such as *P. falciparum*, there is evidence that SPDS can catalyze the formation of both spermidine and spermine from putrescine and spermidine respectively, and a canonical spermine synthase is absent^30^. Further studies in *B. duncani* are needed to determine whether a single enzyme catalyzes the conversion of putrescine into spermidine and spermidine into spermine, or whether this parasite possesses two independent enzymes to synthesize spermidine and spermine from putrescine. Several studies in various organisms including yeast have demonstrated the presence of spermine oxidase that catalyzes the reverse reaction and leads to the conversion of spermine into spermidine ^31^. *In B. duncani*, we identified a spermine oxidase (*BdSMO*) gene (BdWA1_002305) by bioinformatics analysis. The encoded putative enzyme BdSMO shares 21% percent identity with yeast (*Saccharomyces cerevisiae*) FMS1-amine oxidase. Although a recent study showed successful genetic manipulation of *B. duncani* ^32^, the technology for efficient gene knockout in this organism needs to be further optimized. Once optimized, the technology could be used to elucidate the role of this and other putative genes in the biosynthesis of polyamines and polyamine-mediated regulation of protein translation in this parasite.

In this study, we demonstrated that depletion of putrescine severely diminishes the ability of *B. duncani* to complete its life cycle with human RBCs (Fig. 2B). However, putrescine depletion from the growth medium did not result in 100% parasite death. The residual growth seen in media lacking putrescine may be due to low amounts of polyamines present either in serum, a critical supplement for continuous propagation of the parasite in vitro, or in human RBCs. Indeed, when the growth of *B. duncani* in media lacking putrescine was compared between RBCs collected from fresh blood (<30 days) or aged blood (60 and 90 days old), (Fig. S1), residual growth was higher with young RBCs than with aged RBCs. This is consistent with the hypothesis that human RBCs contain polyamines that are depleted over time. However, the polyamine levels even in freshly isolated human RBCs are insufficient to support optimal in vitro parasite growth since the parasites cannot successfully replicate in media lacking polyamines.

Research aimed to elucidate the physiological roles of polyamines in mammalian cells suggested a plethora of functions associated with the alterations in the polyamine pathway including defects in protein translation and cell proliferation. The importance of putrescine salvage in *B. duncani* indicates that either putrescine or its downstream products are essential for parasite growth. Our pharmacological data indicated that spermidine is the key polyamine required for intraerythrocytic proliferation of *B. duncani* (Fig. 4). Spermidine is the substrate for deoxyhypusine synthase, the first enzyme of the two-step hypusination pathway that leads to the formation of a hypusine residue by conjugation of the aminobutyl moiety of spermidine to a specific lysine residue of eukaryotic translation initiation factor 5A (eIF5A) ^23^ (Fig. 6). The hypusination pathway has evolved in eukaryotes and is highly conserved, indicating that the cellular function of eIF5A has been maintained through evolution. For the first time, this study reports that depletion of putrescine affects the hypusination of eIF5A in *B. duncani*. The levels of hypusinated eIF5A (hyp-eIF5A) are significantly reduced in the parasites cultured in putrescine depleted growth medium in comparison to putrescine containing growth medium (Fig. 6F). This finding is consistent with our transcriptomic that confirmed altered expression of genes involved in protein translation.

In summary, our data indicate that unlike *P. falciparum*, *B. duncani* uses a salvage pathway for the biosynthesis of its polyamines. However, in both organisms, the primary function of putrescine and spermine is to produce spermidine for the operation of the translation machinery. Targeting polyamine utilization and the polyamine-eIF5A-hypusine axis present attractive therapeutic opportunities to design novel drugs against human babesiosis and malaria.

## Methods

### Ethics Statement

All animal experiments were approved by the Institutional Animal Care and Use Committees (IACUC) at Yale University (Protocol #2022-11619). Animals were acclimatized for one week after arrival before the start of an experiment. Animals that showed signs of distress or appeared moribund were humanly euthanized using approved protocols.

### In vitro parasite culture of *B. duncani* in human RBCs in different growth media

*B. duncani* parasites were cultured in vitro as previously described ^33, 34^. Parasite growth was monitored and evaluated in the following media: DMEM_b_ (Thermofisher, 11965-092) and DMEM/F12 (Lonza, BE04-687/U1). The media were supplemented with 20% heat-inactivated FBS, 2% 50X HT Media Supplement Hybrid-MaxTM (Sigma, H0137), 1% 200 mM L-Glutamine (Gibco, 25030-081), 1% 100X Penicillin/Streptomycin (Gibco, 15240-062) and 1% 10 mg/mL Gentamicin (Gibco, 15710-072). Parasitemia was monitored by light microscopy examination of Giemsa-stained blood smears.

### Comparison of in vitro growth of *B. duncani* WA-1 in DMEM and DMEM/F12 media

*B. duncani* WA-1 was cultured in vitro in human RBCs in complete DMEM/F12 (cDMEM/F12) medium. The parasitemia was monitored by Giemsa-stained blood smears. An assay to compare the growth of parasites in DMEM medium versus DMEM/F12 medium was set using the parasites grown in cDMEM/F12 medium. Briefly, the parasite culture was washed three times using incomplete DMEM (iDMEM). Next, the parasites were diluted to 0.5% parasitemia (using fresh human RBCs) in iDMEM medium. Subsequently, 2 ml of parasite culture (0.5% parasitemia and 5% HC) was aliquoted into 6 different tubes and centrifuged for 5 min at 1800 rpm at room temperature ^8^. Following this, the supernatant was discarded; cDMEM/F12 was added to three tubes and complete DMEM (cDMEM) was added to the remaining three tubes. The cultures from different tubes were plated in wells of a 12-well plate. A blood smear was prepared from all the 6 wells for Giemsa-staining and the day 0 parasitemia was estimated by counting roughly 2500-3000 RBCs. The cultures were allowed to grow for two days and the parasitemia was measured on day 3^rd^ by counting of Giemsa-stained blood smears. Following this, all the cultures were diluted to 0.5% parasitemia by addition of fresh human A^+^ RBCs and respective growth medium and allowed to grow for two days. Similarly, on day 6^th^ and day 9^th^ the parasitemia was measured and the cultures were diluted to 0.5% parasitemia. From day 9^th^ onwards, the parasites were allowed to grow continuously until day 15^th^ without any dilution. The respective culture media was replaced on day 12^th^ and day 14^th^ and the parasitemia was monitored on day 12^th^ and day 15^th^.

### Comparison of in vitro growth of *B. duncani* WA-1 in cDMEM/F12 and drop-in media

To determine component/s essential for parasite growth present specifically in DMEM/F12 but missing in DMEM, the following media were prepared: cDMEM/F12 (control), cDMEM, cDMEM + missing amino acids (4.45 mg/L of L-Alanine, 7.5 mg/L of L-Asparagine, 6.65 mg/L of L-Aspartic acid, 17.56 mg/L of L-Cysteine, 7.35 mg/L of L-Glutamic acid and 17.27 mg/L of L-Proline), cDMEM + missing vitamins (0.004 mg/L of biotin and 0.68 mg/L of vitamin B12), cDMEM + missing salts (0.0012 mg/L of cupric sulfate, 0.42 mg/L of ferric sulfate, 28.57 mg/L of magnesium chloride and 0.43 mg/L of zinc sulfate), cDMEM + missing lipids (0.044 mg/L of linoleic acid and 0.103 mg/L of lipoic acid), cDMEM + putrescine (0.081 mg/L of putrescine) (Sigma, P5780), cDMEM + missing AAs + Lipids + Putrescine (4.45 mg/L of L-Alanine, 7.5 mg/L of L-Asparagine, 6.65 mg/L of L-Aspartic acid, 17.56 mg/L of L-Cysteine, 7.35 mg/L of L-Glutamic acid, 17.27 mg/L of L-Proline, 0.044 mg/L of linoleic acid, 0.103 mg/L of lipoic acid and 0.081 mg/L of putrescine) and cDMEM + missing lipids and putrescine (0.044 mg/L of linoleic acid, 0.103 mg/L of lipoic acid and 0.081 mg/L of putrescine). For this growth assay, the *B. duncani* WA-1 parasites were grown in cDMEM/F12 to 10% parasitemia. The parasites were washed three times in iDMEM and diluted to 0.5% parasitemia (5% HC). Two milliliter cultures (0.5% parasitemia and 5% HC) were aliquoted in different tubes (3 replicate tubes per media to be tested). Following this, the tubes were centrifuged to pellet the culture and the supernatants were discarded. Then, 2 ml of the respective media was added to three different tubes (replicates) and plated in the wells of a 12-well plate. A blood smear was prepared from all the wells for Giemsa-staining and the day 0 parasitemia was estimated by counting roughly 2500-3000 RBCs. The cultures were allowed to grow for two days and the parasitemia was measured on day 3^rd^ by counting of Giemsa-stained blood smears. Following this, all the cultures were diluted to 0.5% parasitemia by addition of fresh human A^+^ RBCs and respective growth medium and allowed to grow for two days. Similarly, on day 6^th^ and day 9^th^ the parasitemia was measured and the cultures were diluted to 0.5% parasitemia. From day 9^th^ onwards, the parasites were allowed to grow continuously for up to day 15^th^ without any dilution. The respective culture media was replaced on day 12^th^ and day 14^th^ and the parasitemia was monitored on day 12^th^ and day 15^th^.

### Comparison of in vitro growth of *B. duncani* WA-1 in cDMEM/F12 and drop-out media

To determine which amino acid or lipids or putrescine is essential for parasite growth present specifically in DMEM/F12 but missing in DMEM, the following media were prepared: cDMEM/F12 (control), cDMEM, cDMEM +All -LNA (DMEM supplemented with all missing components except linoleic acid), cDMEM +All -LA (DMEM + All except lipoic acid), cDMEM +All -Put (DMEM + All except putrescine), cDMEM +All -Ala (DMEM + All except L-alanine), cDMEM +All -Asn (DMEM +All except L-asparagine), cDMEM +All -Pro (DMEM + All except L-proline), cDMEM +All -Cys (DMEM + All except L-cysteine) and cDMEM +All -Glu (DMEM + All except L-glutamic acid). For setting up this growth assay, the *B. duncani* WA-1 parasites were grown in cDMEM/F12 to 10% parasitemia. The parasites were washed three times in iDMEM and diluted to 0.5% parasitemia (5% HC). Two milliliter cultures (0.5% parasitemia and 5% HC) were aliquoted in different tubes (3 replicate tubes per media to be tested). Following this, the tubes were centrifuged to pellet the culture and the supernatants were discarded. Subsequently 2 ml of the respective media was added to three different tubes (replicates) and plated in the wells of a 12-well plate. A blood smear was prepared from all the wells for Giemsa-staining and the day 0 parasitemia was estimated by counting roughly 2500-3000 RBCs. The cultures were allowed to grow for two days and the parasitemia was measured on day 3^rd^ by counting of Giemsa-stained blood smears. Following this, all the cultures were diluted to 0.5% parasitemia by addition of fresh human A^+^ RBCs and respective growth medium and allowed to grow for two days. Similarly, on day 6^th^ and day 9^th^ the parasitemia was measured and the cultures were diluted to 0.5% parasitemia. From day 9^th^ onwards, the parasites were allowed to grow continuously for up to day 15^th^ without any dilution. The respective culture media was replaced on day 12^th^ and day 14^th^ and the parasitemia was monitored on day 12^th^ and day 15^th^.

### Ornithine supplementation growth assay in putrescine depleted *B. duncani* WA-1 parasites

To test whether the growth defect of parasites grown in putrescine depleted media can be rescued by addition of ornithine (the precursor of putrescine), *B. duncani* parasites were grown in putrescine containing medium (cDMEM/F12) to 10% parasitemia. The parasites were washed three times in iDMEM, and the cultures was diluted to 0.5 % parasitemia (5% HC) in iDMEM. One milliliter cultures (0.5% parasitemia and 5% HC in iDMEM medium) were aliquoted in 15 different microcentrifuge tubes (3 replicate tubes per experimental condition). The cultures were centrifuged for 5 min at 1800 rpm at RT and the supernatants were discarded. To first set of three microcentrifuge tubes (triplicates), 1 ml of putrescine containing medium (cDMEM/F12), was added (positive control). To the remaining 12 tubes, 1 ml each of putrescine depleted medium (cDMEM + All except putrescine) was added. To a set of 3 tubes (0.1 mg/ml L-Ornithine), 1 μl of 100 mg/ml of L-Ornithine was added. To a second set of 3 tubes (0.5 mg/ml Ornithine), 5 μl of 100 mg/ml of L-Ornithine was added. To the third set of 3 tubes (1 mg/ml Ornithine), 10 μl of 100 mg/ml of L-Ornithine was added. The cultures from all the tubes were seeded in the wells of a 24-well plate. Day 0 smears were prepared and the parasitemia was estimated. The cultures were allowed to grow continuously for 6 days, with media change (respective media as listed above) on day 3^rd^. The final parasitemia on day 6 was monitored by examination of Giemsa-stained blood smears.

### Spermidine and spermine supplementation growth assay in putrescine depleted *B. duncani* parasites

To test whether spermidine and spermine supplementation can rescue the growth defect of parasites grown in putrescine depleted media, *B. duncani* parasites were grown in putrescine containing medium (cDMEM/F12) to 10% parasitemia. The parasites were washed three times in iDMEM, and the culture was diluted to 0.5 % parasitemia (5% HC) in iDMEM. One milliliter cultures (0.5% parasitemia and 5% HC in iDMEM medium) were aliquoted in 24 different microcentrifuge tubes (3 replicate tubes per experimental condition). The cultures were centrifuged for 5 min at 1800 rpm at RT and the supernatants were discarded. To first set of three microcentrifuge tubes (triplicates), 1 ml of putrescine containing medium (cDMEM/F12), was added (positive control). To the remaining 21 tubes, 1 ml of putrescine depleted medium (cDMEM + All except putrescine) was added. Out of the 21 tubes, 3 were kept as is, whereas the remaining tubes were supplemented with either spermine or spermidine. To a set of 3 tubes (0.1 mg/ml spermidine), 1μl of 100 mg/ml of spermidine (Sigma, S0266) was added. Similarly, to the 2^nd^ set of 3 tubes (0.5 mg/ml spermidine) and 3^rd^ set of 3 tubes (1 mg/ml spermidine), 5μl of 100 mg/ml and 10μl of 100 mg/ml of spermidine was added respectively. To the 4^th^ set of 3 tubes (0.1 mg/ml spermine), 2 μl of 50 mg/ml of spermine (Sigma, S4264) was added. Similarly, to the 6^th^ set of 3 tubes (0.5 mg/ml spermine) and 7^th^ set of 3 tubes (1 mg/ml spermine), 10μl of 50 mg/ml and 20μl of 50 mg/ml of spermine was added respectively. The cultures from all the tubes were seeded in the wells of a 24-well plate. Day 0 smears were prepared and the parasitemia was estimated. The cultures were allowed to grow continuously for 6 days, with media change (respective media as listed above) on day 3^rd^. The final parasitemia on day 6 was monitored by examination of Giemsa-stained blood smears.

### In vitro susceptibility of *B. duncani* WA-1 to DFMO and DFMA

The susceptibility of the intraerythrocytic development cycle of *B. duncani* WA-1 cultured in putrescine containing or depleted media to DL-α-difluoromethylornithine (DFMO) (Sigma, D193) and DL-α-difluoromethylarginine (DFMA) (Cayman chemical, 16415) was determined by incubating the parasite cultures (0.5% parasitemia, 5% HC in either putrescine containing or depleted medium) to different concentrations of these drugs (5 mM, 2.5 mM, 1.3 mM, and 0.6 mM of DFMO; 250 mM, 125 mM, 62.5 mM, and 31.3 mM of DFMA). The untreated parasites from putrescine containing (Plus Putrescine) or depleted media (Minus Putrescine) were used as no drug controls. Uninfected RBCs (5% HC) were used as negative control. The assays were performed in triplicates in a 96-well plate in 200 μl volume and maintained for 60 h at 37°C in an incubator with a mixture of 2% O_2_, 5% CO_2_, and 93% N_2_. Subsequently, 100 μl of the culture per well of the 96 well plate was used in a SYBR Green I assay. Briefly, 100 μl of the culture from the 96-well assay plate was transferred to the Costar 96-well black plate and 100 μl of SYBR Green I lysis buffer (20 mM Tris (pH 7.4), 5 mM EDTA, 0.008% saponin, 0.08% Triton X-100, and 1X SYBR Green I (Molecular Probes, 10,000X solution in DMSO)) was added to the wells containing the parasite culture. The plate was incubated at 37°C in the dark for 1 h and subsequently read on a BioTek SynergyMX fluorescence plate reader with an excitation of 497 nm and emission of 520 nm. Three independent experiments were performed in biological triplicates and the data was analyzed using GraphPad Prism (Version 9.4.1).

### Perturbation of the intraerythrocytic cycle of *B. duncani* WA-1 with 4-Methylcyclohexylamine (4-MCHA), MDL 72527 and GC7

*B. duncani* WA-1 parasites were cultured in in vitro in cDMEM/F12 medium or cDMEM/F12 supplemented with either 1 mg/ml of putrescine or spermidine or spermine. The cultures maintained in the above mentioned media were diluted to 0.5% parasitemia (5%HC) and treated with decreasing concentrations (2-fold dilution starting from 10mM (4-MCHA), 1mM (MDL 72527) and 0.4mM (GC7)) of different drugs, including: 4-MCHA (Sigma, 177466), MDL 72527 (Sigma, M2949) and GC7 (Santa Cruz, sc-396111) for 60 h in a 96-well plate. The parasite viability was determined by the SYBR green I method (mentioned above). Similarly, *B. duncani* parasites were grown in putrescine depleted medium (DMEM +All -Putrescine) or putrescine depleted medium supplemented with 0.1 mg/ml, 0.5 mg/ml and 1 mg/ml of putrescine or putrescine depleted medium supplemented with 1 mg/ml of spermidine or spermine. The parasites grown in the above mentioned different growth media were diluted to 0.5% parasitemia (5%HC) and treated with decreasing concentrations (2-fold dilution starting from 10mM (4-MCHA), 1mM (MDL 72527) and 0.4mM (GC7)) of different drugs, including: 4-MCHA, MDL and GC7 for 60 h in a 96-well plate. The parasite viability was determined by the SYBR green I method (mentioned above). Three independent experiments were performed in biological triplicates and the data was analyzed using GraphPad Prism (Version 9.4.1).

### Immunoblot analysis

*B. duncani* parasite lysate from parasites grown in different media (putrescine containing, putrescine depleted, putrescine depleted media supplemented with spermidine or spermine supplemented) were collected. Briefly, the *B. duncani* WA-1 infected RBC pellets was treated with 0.1% saponin (Sigma, SAE0073) to lyse the RBCs and the parasite pellet was washed three times with 1X PBS. Subsequently, the parasite pellets were resuspended in 1X Laemmli buffer (Bio-Rad, 161-0747) containing 5% β-mercaptoethanol (Bio-Rad, 1610710), boiled for 5 minutes at 95°C and loaded onto Mini-Protean TGX Stain Free Gels 4-20% (Bio-Rad, P4568096). The gels were transferred onto 0.45 μm nitrocellulose membranes (Bio-Rad, 1620115) and blocked in 5% skimmed milk for 2 H at RT. Following this, the membranes were exposed to either anti-eIF5A rabbit polyclonal antibody (1:500) (Abcam, ab137561) or anti-hypusine rabbit polyclonal antibody (1:500) (Sigma, ABS1064-I) for overnight at 4°C. As a loading control, anti-Bdhsp70-2 rabbit polyclonal antisera was used. Following the primary antibody incubation overnight, the membranes were washed 3 times in 1X PBS containing 0.1% Tween-20 (Thermofisher Scientific, 85114) (PBS-T) and 2 times in 1X PBS. The membranes were then incubated with goat anti-rabbit IgG (H+L) HRP conjugated secondary antibody (1:5000) (Thermofisher Scientific, 31460) for 1 hour at RT. Following this, the membranes were washed 3X in PBS-T and 2X in 1X PBS, and the signals were developed using Super signalTM West Pico PLUS chemiluminescent substrate (Thermo Scientific, 34577). The blot was imaged using LI-COR Odyssey-Fc imaging system. Signal from anti-Bdhsp70-2 rabbit serum was used for normalization.

### Sample collection for RNA sequencing

*B. duncani* parasites were cultured in vitro in putrescine containing growth medium to 10% parasitemia. Following this, the cultures were collected in microcentrifuge tubes and washed 5 times with incomplete DMEM medium. The cultures were then diluted to 1% parasitemia using fresh human RBCs and either putrescine containing or depleted growth media. A total of 25 ml culture (1% parasitemia and 5% HC) in duplicate plates in either putrescine containing, or depleted growth medium were seeded and allowed to grow for 24 H. After this, the cultures were harvested in different tubes and the pellets were resuspended in 5ml trizol (T9424, Sigma) each. Following this, 1 ml chloroform was added to each tube, mixed vigorously for 5 min, and incubated for 10 min at RT. Following this, the tubes were centrifuged at 13,000 rpm for 20 min. The aqueous layer was collected, and equal volume of isopropanol was added, mixed, and incubated for 10 min at RT. Following this, the tubes were centrifuged at 13,000 rpm for 10 min and supernatant was discarded. The pellets were washed twice in freshly prepared 70% ethanol and allowed to air dry after the final wash. The pellets were resuspended in 25 μl of DEPC-treated water. The concentration and quality of RNA was estimated using nanodrop (Synergy HTX multimode reader, Agilent) and Illumina RNA sequencing was performed on the samples. Total RNA was then treated with DNA-free DNA removal kit (ThermoFisher; AM1906) followed by mRNA purification using NEBNext Poly(A) mRNA Magnetic Isolation Module (NEB, E7490S). RNA-seq library was constructed using NEBNext Ultra II RNA-library preparation kit (NEB, E7770S) according to the manufacturer’s instructions. The libraries were amplified for 15 PCR cycles (45s at 98°C followed by 15 cycles of [15s at 98°C, 30s at 55°C, 30s at 62°C], 5 min 62°C). Libraries were subjected to at 150 bp paired end sequencing on the Illumina Novaseq platform.

### Flow cytometry to estimate reactive oxygen species (ROS) in *B. duncani* under putrescine depleted state using dihydrorhodamine 123 dye

The *B. duncani* WA-1 cultures were initiated at 0.5% parasitemia in different growth media, including putrescine containing or depleted media, or putrescine depleted medium supplemented with 1 mg/ml spermidine or 1 mg/ml spermine in 1 ml volume. All the cultures were allowed to grow for two days. Parallelly, another set of parasites grown in putrescine containing media were treated with different concentrations of artemisinin (20nM, 50nM, 0.5μM or 1μM) for two days. On day 3^rd^, all the cultures were washed twice with 1X PBS and resuspended in 1ml of 1XPBS. 0.25 μg/ml of dihydrorhodamine 123 (Sigma, D1054) and Hoechst H33342 (1:1000) were added to all cultures and allowed to incubate for 30 min at 37°C in dark. Following this, the cultures were washed twice with 1X PBS to remove the dyes and resuspended in 1ml of 1X PBS in dark. Unstained *B. duncani* WA-1 culture and uninfected human RBCs were used as negative controls. The cells were analyzed in a flow cytometer (BD LSRII). The cells positive for rhodamine (formed by breakdown of dihydrorhodamine 123 through action of reactive oxygen species) and Hoechst were estimated. The number of rhodamine positive *B. duncani* WA-1 infected RBCs in putrescine depleted medium was used for normalization. The data was plotted on GraphPad prism and the statistical significance of differences between parasite growth in different growth media was estimated by multiple t-tests.

### In vitro parasite culture of *P. falciparum* 3D7 in human RBCs

*P. falciparum* 3D7 blood-stage cultures were maintained at 4% hematocrit in human A^+^ erythrocytes in Rosewell Park Memorial Institute (RPMI) 1640/Hepes medium (Gibco, 11875119) supplemented with 0.5% Albumax II (Gibco, 11021037), HT media supplement (Sigma, H0137), and gentamicin (Thermofisher Scientific, 15710-072) under mixed gas (5% O_2_, 5% CO_2_, and 90% N_2_). Parasitemia was monitored by light microscopy examination of Giemsa-stained blood smears.

## Author Contributions

P.S.: Investigation, methodology, analysis, visualization, writing the original draft, review and editing. C.B.M.: Conceptualization, supervision, funding acquisition, project administration, writing original draft, review, and editing. All authors have read and agreed to the published version of the manuscript.

## Acknowledgements

We thank Meenal Chand for help with the immunofluorescence assays and Joseph Gennaro for commenting on the manuscript.

---

**Figure S1.**
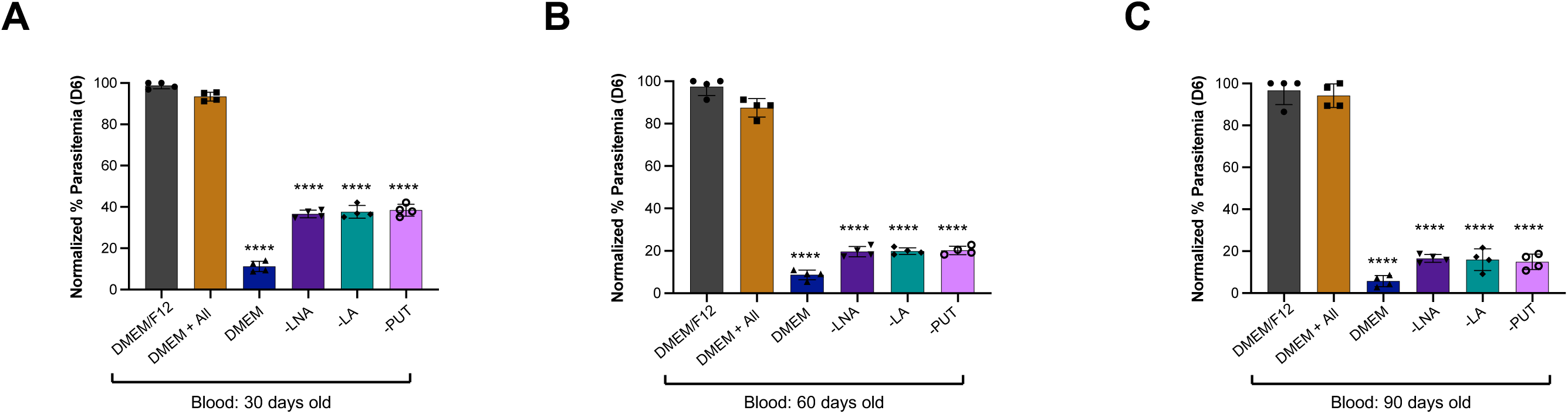
Growth defect in *B. duncani* upon removal of putrescine (PUT), linoleic acid ^35^ and lipoic acid (LA) is more pronounced in aged blood. **A.** Comparison of *B. duncani* growth in DMEM/F12, DMEM, DMEM containing all the missing components (DMEM + All) or DMEM+All depleted in individual components (linoleic acid (LNA), lipoic acid (LA), putrescine (PUT)), in 30 days old human A^+^ RBCs over 6 days. The parasitemia was monitored at the end of day 6. A total of 3000-3500 RBCs were counted. The parasite growth in DMEM/F12 medium was used for normalization. The normalized percent parasitemia at the end of day 6 is depicted in the graph. Data presented as mean ± SD of two independent experiments performed in biological duplicates. Statistical significance of differences between parasite growth in DMEM/F12 and other media was estimated by Welch’s t-test. **** indicates significant P values <0.0001. **B.** Comparison of *B. duncani* growth in DMEM/F12, DMEM, DMEM (DMEM + All) or DMEM+All depleted in individual components (linoleic acid (LNA), lipoic acid (LA), putrescine (PUT)), in 60 days old human A^+^ RBCs over 6 days. The parasitemia was monitored at the end of day 6. A total of 3000-3500 RBCs were counted. The parasite growth in DMEM/F12 medium was used for normalization. The normalized percent parasitemia at the end of day 6 is depicted in the graph. Data presented as mean ± SD of two independent experiments performed in biological duplicates. Statistical significance of differences between parasite growth in DMEM/F12 and other media was estimated by Welch’s t-test. **** indicates significant P values <0.0001. **C.** Comparison of *B. duncani* growth in DMEM/F12, DMEM, DMEM (DMEM + All) or DMEM+All depleted in individual components (linoleic acid (LNA), lipoic acid (LA), putrescine (PUT)), in 90 days old human A^+^ RBCs over 6 days. The parasitemia was monitored at the end of day 6. A total of 3000-3500 RBCs were counted. The parasite growth in DMEM/F12 medium was used for normalization. The normalized percent parasitemia at the end of day 6 is depicted in the graph. Data presented as mean ± SD of two independent experiments performed in biological duplicates. Statistical significance of differences between parasite growth in DMEM/F12 and other media was estimated by Welch’s t-test. **** indicates significant P values <0.0001.

**Figure S2.**
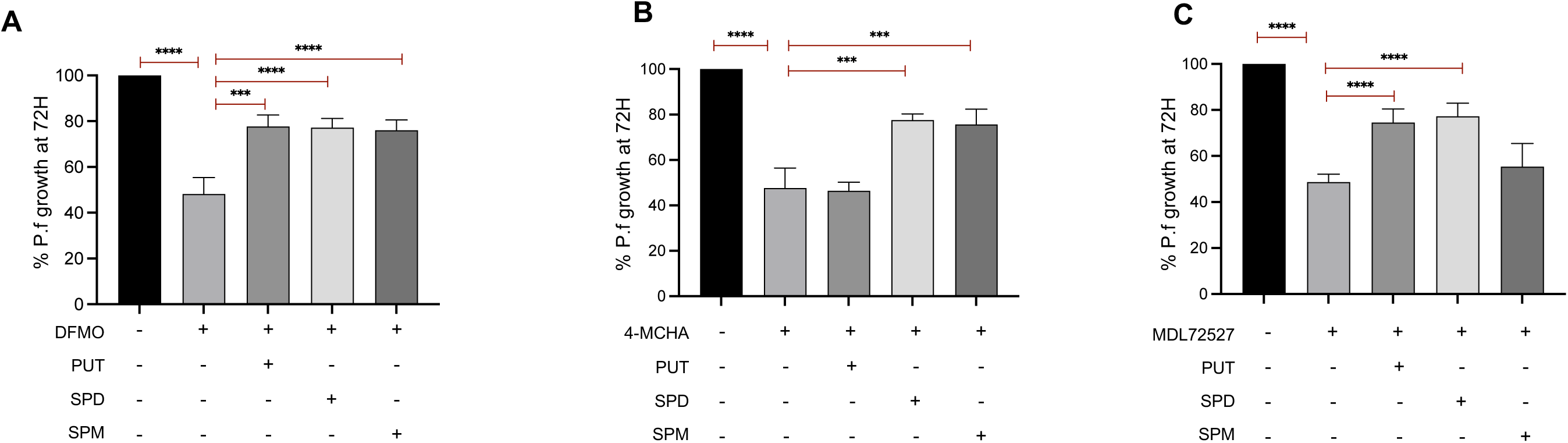
Inhibition of polyamine biosynthesis pathway in *P. falciparum* 3D7 can be reversed by the addition of polyamines. **A.** Inhibition of *B. duncani* growth with IC_50_ concentration (1mM) of DL-α-difluoromethylornithine (DFMO) in the presence or absence of different polyamines (putrescine (PUT), spermidine (SPD) and spermine (SPM)). Data presented as mean ± SD of two independent experiments performed in biological triplicates. Statistical significance of differences was estimated by Welch’s t-test. **** indicates significant P values <0.0001. **B.** Inhibition of *B. duncani* growth with IC_50_ concentration (90 μM) of 4-MCHA in the presence or absence of different polyamines (putrescine (PUT), spermidine (SPD) and spermine (SPM)). Data presented as mean ± SD of two independent experiments performed in biological triplicates. Statistical significance of differences was estimated by Welch’s t-test. *** indicates significant P values <0.001, **** indicates significant P values <0.0001. **C.** Inhibition of *B. duncani* growth with IC_50_ concentration (87 μM) of MDL 72527 in the presence or absence of different polyamines (putrescine (PUT), spermidine (SPD) and spermine (SPM)). Data presented as mean ± SD of two independent experiments performed in biological triplicates. Statistical significance of differences was estimated by Welch’s t-test. *** indicates significant P values <0.001, **** indicates significant P values <0.0001.

## Notes

**Conflict of interest:** None of the named authors have any conflict of interest, financial or otherwise.

Funding: This work was supported by the National Institutes of Health AI153100. CBM research is also supported by NIH grants AI138139, AI152220, AI123321 and AI136118, the Steven and Alexandra Cohen Foundation [Lyme 62 2020 to CBM], and the Global Lyme Alliance.

### Competing Interest Statement

The authors have declared no competing interest.

